# An exact, unifying framework for region-based association testing in family-based designs, including higher criticism approaches, SKATs, multivariate and burden tests

**DOI:** 10.1101/815290

**Authors:** Julian Hecker, F. William Townes, Priyadarshini Kachroo, Jessica Lasky-Su, John Ziniti, Michael H. Cho, Scott T. Weiss, Nan M. Laird, Christoph Lange

## Abstract

Analysis of rare variants in family-based studies remains a challenge. To perform a region/set-based association analysis of rare variants in family-based studies, we propose a general methodological framework that integrates higher criticism, maximum, SKATs, and burden approaches into the family-based association testing (FBAT) framework. Using the haplotype algorithm for FBATs to compute the conditional genotype distribution under the null hypothesis of Mendelian transmissions, virtually any association test statistics can be implemented in our approach and simulation-based or exact p-values can be computed without the need for asymptotic settings. Using simulations, we compare the features of the proposed test statistics in our framework with the existing region-based methodology for family-based studies under various scenarios. The tests of our framework outperform the existing approaches. We provide general guidelines for which scenarios, e.g., sparseness of the signals or local LD structure, which test statistic will have distinct power advantages over the others. We also illustrate our approach in an application to a whole-genome sequencing dataset with 897 asthmatic trios.

## Introduction

In family-based association studies, the concept of Mendelian transmissions can be utilized to construct association tests that are robust against genetic confounding (Transmission Disequilibrium Tests (TDTs) ^1^ or Family-based Association Tests (FBATs) ^2^). This feature of family-based association tests was fundamental to establish them as a popular tool in association mapping since the days of candidate gene studies. The robustness of the approach largely out-weighted the requirement of family data, i.e., having to recruit additional related study subjects, and reduced statistical power compared to population-based cohorts with the same sample size. Testing strategies have been proposed that allowed the incorporation of the association information at the population-level in family-based designs without compromising the robustness of the test statistic ^3–7^.

As genome-wide association studies (GWAS) became a standard research tool, and the multiple testing problem had to be addressed at a genome-wide level, researchers started to emphasize statistical power in their choice for study designs. Power and study design considerations substantially contributed to making population-based designs approaches the most popular choice in association analysis.

Now, as whole-genome sequencing (WGS) studies are replacing chip-based GWAS, region-based rare variant analysis approaches have moved to the center of the statistical methodology development, as, even for very large sample sizes, the power of single locus association analysis will be too small when the minor allele is rare. Region-based approaches are motivated by the idea that, if we can combine “association signals” across a pre-defined region in a suitable way, a stronger genetic signal could be assessed by a suitable test statistic, and the resulting region-based association test would have increased statistical power. At the same time, the multiple testing problem could become less severe as fewer association tests are computed. The major statistical challenges for region-based tests are to identify suitable ways to combine the genetic information across the pre-specified region, to incorporate/estimate the correlation between the selected rare variants in the region-based test statistic, and to select a suitable test statistic.

For population-based designs, based on assumptions about the alternative hypothesis/distribution of disease susceptibility loci (DSLs) and their effect directions, numerous region-based tests have been proposed, e.g., burden tests and variance component tests ^8,9^. However, population stratification is a potential problem in population-based designs that can be even more severe in the analysis of rare variants.

For family-based designs, the popular burden tests and the SKAT approach have been translated to the FBAT framework ^10,11^. As with their population-based equivalents, these two approaches estimate the correlation between the genetic loci empirically. This, especially for rare variant data, can be problematic, as the rare variant allele counts are small, and the empirical estimates can be affected by numerical instabilities. Furthermore, the application of asymptotic theory may not provide accurate results here. Other recent approaches are based on mixed models and can, theoretically, analyze unrelated and related samples ^12^.

As an other alternative to the transmission-based FBAT approach, methods that compare the allele frequencies between affected and unaffected individuals within families have been suggested, e.g., Generalized Disequilibrium Test (GDT) for single variant analysis ^13^. As they can incorporate all available phenotypic and genetic information within one pedigree, they can be more powerful than the corresponding FBAT analysis. However, they require the assumption that the allele frequencies and the genetic variance for all members of a pedigree are equal under the null hypothesis. This assumption can be violated in the presence of population substructure within the founders of the families or when there are departures from Hardy Weinberg equilibrium ^14^. None of these assumptions is required for the FBAT approach to be valid. For region-based analysis, the Rare-Variant Generalized Disequilibrium Test (RV-GDT) extension has been proposed ^15^.

In this communication, we will exclusively focus on transmission-based analysis approaches to construct association tests that are robust against confounding. We propose a general framework for region-based rare variant analysis in extended pedigree/nuclear families that is based on the FBAT approach. Multiple offspring per family may be available, founder/phase information can be missing, and phenotypes can be dichotomous or quantitative. In contrast to previous approaches for region-based analysis in population- and family-based designs, the joint distribution of the rare variants in the region is obtained analytically under the null hypothesis, conditioning on the sufficient statistic, using the haplotype algorithm for FBATs^16,17^. Based on the conditional genotype distribution, it is straightforward to implement region-based FBATs, e.g., multivariate tests, burden tests, SKAT ^9,11^, and higher criticism approaches ^18–20^. As the joint distribution of the rare variants is obtained analytically and can efficiently be sampled from, the significance of the test statistics in our framework can be obtained either by simulations or the construction of the exact distribution^21^. This flexibility of our approach enables the implementation of virtually any region/set-based test without the need for any asymptotic assumptions or approximations. We illustrate the implementation of higher criticism approaches, maximum statistics, SKATs and burden tests in our framework.

For different scenarios, e.g., regions with sparse signals, varying local Linkage Disequilibrium (LD) structure, we compare our proposed FBAT framework to existing methodology, using extensive simulation studies. Our simulation results support our theoretical considerations that our testing framework provides a substantial improvement over the existing methodology in terms of statistical power and robustness against population substructure. We also develop general recommendations for which choice of the test statistic is preferable for which scenario. Furthermore, we also applied our methodology framework to a whole-genome sequencing study for childhood asthma with 897 trios.

## Methods

In a family-based WGS association study, genotype data for rare variants are available for a set of marker loci that are in close physical proximity and define a genomic segment that is suitable for region-based association analysis. The genotype information may be available for multiple offspring as well as for the parents. For the *i*-th nuclear family, we introduce the *p* × *n*_*i*_ genotype matrix *X*_*i*_ and the *n*_*i*_ dimensional phenotype vector *T*_*i*_, where *n*_*i*_ denotes the number of offspring in the i-th nuclear family, and *p* denotes the number of variants in the analysis region. We regard *X*_*i*_ as random while *T*_*i*_ is fixed in the FBAT approach. Below, we propose a possible set of test statistics that can capture the potential association between the offspring genotype data and the phenotypes under various conditions.

### Simulation-based significance testing

For each region, using the haplotype algorithm for FBAT ^16,17^, we derive the conditional distribution of offspring genotypes *X*_*i*_ in the *i*-th nuclear family under the null hypothesis, given the sufficient statistic *S*_*i*_ for the possible missing founder genotypes ^22^. The sufficient statistic approach utilizes parental genotypes if they are available. The knowledge about the conditional genotype distribution allows constructing association tests that are robust against population stratification and admixture. Based on this conditional genotype distribution, it is straightforward to compute the first two moments of commonly used test statistics under the null hypothesis of no association and derive the asymptotic distribution. However, as the analysis of rare variant data leads to scenarios where the application of asymptotic theory does not provide reliable approximations, simulation-based or even exact p-values ^21^ are preferred. Here, we propose to evaluate association p-values based on a sufficiently large number of simulated draws from the null distribution. This procedure can be combined with adaptive permutation/simulation-based p-value techniques. In this context, we recommend using stopping rules that are nearly optimal in terms of the required number of simulations ^23^.

This approach, therefore, also allows our testing framework to incorporate test statistics where the (asymptotic) distribution is intractable or cumbersome, e.g., maximum statistics or higher criticism approaches.

### Test statistics

All test statistics under consideration are based on the following two objects. For the i-th family, we define the *p*-dimensional vector of Mendelian residuals *U*_*i*_ = (*X*_*i*_ − *E*[*X*_*i*_|*S*_*i*_])*T*_*i*_. Also, we define the corresponding *p* × *p* variance matrix *V*_*i*_ = *Var*(*U*_*i*_|*S*_*i*_, *T*_*i*_). For both objects, the moments are computed under the null hypothesis, based on the sufficient statistic *S*_*i*_.

### Burden-based approaches

Burden-type FBATs can be implemented by specifying a *p*-dimensional weight vector *W* that collapses/summarizes the rare variant information of the region into a single scalar value. The specification of the weight vector *W* requires assumptions about the effect direction and its effect size. In this context, the contribution to the FBAT statistic of the *i*-th family is then given by

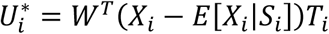

The corresponding FBAT-statistic for the simulation-based testing is computed by 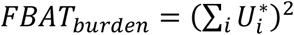. We note that for this burden test, it would be possible as well to compute an asymptotic p-value by also computing/estimating the corresponding variance.

### Variance component/SKAT approaches

As an alternative to burden/collapsing association tests, SKAT/variance-component based region tests have been developed for rare variant data ^9^. They have the advantage that they do not require any assumptions about the effect configuration at the rare variant loci under the alternative hypothesis, but they are not as powerful as burden/collapsing approaches if one is certain about the alternative hypothesis. We define the general statistic

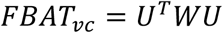

where *U* = ∑_*i*_ *U*_*i*_ and *W* is a fixed *p* × *p* weight matrix. While, in the scenario of affected offspring trios and a diagonal weight matrix *W*, this test statistic equals the FB-SKAT statistic ^11^, Ionita-Laza et al. assess significance based on asymptotic results that require the empirical estimation of the variance/covariance matrix for the rare variants, which, given the sparseness of rare variant data, can become problematic. In our framework, the p-value of the test statistic is obtained based on simulations from the conditional genotype distribution.

If we set *W* = *V*^−1^, where *V* = ∑_*i*_ *V*_*i*_, we obtain the multivariate FBAT ^24^. The multivariate FBAT was designed for common variants, and the asymptotic p-values also require an empirical estimate of the correlation matrix. Again, for rare variants, this can lead to unreliable results, making the implementation of the multivariate FBAT in our proposed framework preferable.

### Higher criticism and maximum statistic

Besides the commonly used burden and variance component approaches, we introduce the higher criticism and maximum statistic for region-based analysis in family-based studies. Both approaches are designed to identify sparse alternatives and have been introduced to genetic association studies of unrelated individuals recently ^18,20^.

Define the normalized residuals 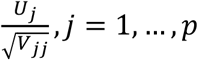 and denote the corresponding association p-value based on the asymptotic marginal normal distribution by *q*_*j*_. Based on the available amount of information per variant, e.g., the number of informative transmissions/families, we restrict the set of variants to a subset of variants where the marginal variance is large enough (e.g., we require at least 5 informative nuclear families). Denote the number of variants in this subset by *p*′. Given the ordered p-values *q*_(1)_ ≤ *q*_(2)_ ≤ … ≤ *q*_(*p*′)_, we define the HC statistic as

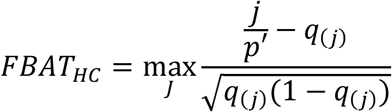

Here, the index set *J* can be >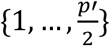 or {1, … , *p*′}, depending on assumptions about the underlying genetic architecture.

It is important to note that, while *FBAT*_*HC*_ contains a transformation based on the single variant asymptotic distribution, the assessment of its significance based on simulations from the conditional distribution remains a valid approach regardless of whether the assumptions that motivated the transformation hold.

The second approach to detect spare signals in the tested genomic region is the MAX statistic which is simply defined as

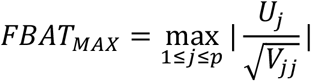

The theory related to the higher criticism/max statistic in the setting of unrelated case-control data and sparse signals developed in Mukherjee et al. ^20^ can be transferred to family-based studies/FBATS. In Appendix A, we derive how the theory in Mukherjee et al. ^20^ can be applied to the FBAT framework in the scenario of affected offspring trios. The corresponding optimality results for sparse signal scenarios motivate the application of these test statistics to rare variants in sequencing studies.

Finally, we note that it is, of course, possible to set up an omnibus statistic that is based on the maximum of multiple test statistics described above.

## Results

In this section, we describe the results of two simulation studies and an analysis of a whole-genome sequencing study for childhood-asthma with 897 affected offspring trios.

### Simulation studies

We studied the performance of our proposed test statistics in two extensive simulation studies. In both studies, we compared the Type I error and power with the existing methodology for family-based region association analysis. We restricted all simulations to the scenario of trios with an affected offspring. However, it is important to note that our framework can be applied to any nuclear family and phenotype distribution. For the test statistics *FBAT*_*burden*_ and *FBAT*_*vc*_, we applied uniform weights. In the following, we will denote the test statistics *FBAT*_*burden*_, *FBAT*_*vc*_, *FBAT*_*HC*_, and *FBAT*_*MAX*_ by Burden, SKAT, HC, and MAX.

#### Genetic regions with unphased data

We extracted haplotypes for the CEU and the GBR subpopulations from the 1000 Genomes Project ^25^, consisting of 30 and 50 consecutive rare variants with a minor allele frequency (MAF) below 3%. Based on these haplotypes, we generated genotype data for trios using Mendelian transmissions. Using a standard logistic disease model with a disease prevalence of ≈ 10%, we simulated offspring affection status and collected *n* = 1,000 affected offspring trios. This simulation study is similar to the simulation studies described in the existing literature ^15,26^.

We compared our test statistics with the GTDT ^26^ and the RV-GDT ^15^. The GTDT ^26^ offers five different test statistics for region-based affected offspring trio analysis, designed for different modes of inheritance. The test statistics require phased haplotype data. If the phase information is not available, this information is reconstructed up to small uncertainties. We considered the test statistics GTDT-AD, GTDT-DOM, and GTDT-CH in our study. The RV-GDT ^15^ describes a generalization of the single variant GDT ^13^ for multiple variants in a genetic region. The RV-GDT can be applied to arbitrary pedigrees where affected, and unaffected samples are collected; members can be missing. The test statistic compares the genotype counts between affected and unaffected members and corrects for the relatedness using the estimated/reported kinship coefficients. We note that this implies that the phenotype information for parents must be available, whereas the classical TDT/FBAT test for offspring trios does not require this information. For comparison, we included the test MAX-BF that tests if at least one single variant FBAT statistic reached the Bonferroni-corrected significance level corresponding to the number of variants *p* in the region. The corresponding single variant p-values were evaluated using asymptotic theory due to computational reasons.

To check the Type I error rates and the robustness against population stratification; we considered a null hypothesis simulation where no genetic variant is associated with the affection status and three different population admixture scenarios (Table 1). In these admixture scenarios, we generated one fixed parent based on the CEU haplotypes and the other parent based on the GBR haplotypes. The affection status of the parents differed across the three admixture scenarios. For the power analysis, we simulated six different scenarios where the number of causal variants and corresponding effect sizes differ (Table 2). In scenario 5, we picked very rare and independent causal variants with a MAF below 1%, and in scenario 6 we chose causal variants that are in strong LD with multiple other variants. All results are based on 1,000 replicates.

**Table 1.** Type I errors at a significance level of 5% for the FBAT, GTDT and RV-GDT statistics. We considered four scenarios, separately for *p* = 30 and *p* = 50 variants. All results based on 1,000 replicates. null: no association between genetic variants and phenotype, no population admixture adm1: 1 parent generated from CEU haplotypes, 1 parent generated from GBR haplotypes. Parents unaffected, offspring affected. adm2: 1 parent generated from CEU haplotypes, 1 parent generated from GBR haplotypes. CEU parent affected, GBR parent unaffected, offspring affected. adm3: 1 parent generated from CEU haplotypes, 1 parent generated from GBR haplotypes. CEU parent unaffected, GBR parent affected, offspring affected.

|  | FBAT |  |  |  |  | GTD |  |  | RV-GDT |
| --- | --- | --- | --- | --- | --- | --- | --- | --- | --- |
|  | Burden | SKAT | MAX | HC | MAX-BF | GTD-AD | GTD-DOM | GTD-CH | RV-GDT |
| **$p = 30$** |  |  |  |  |  |  |  |  |  |
| null | 4.7% | 5.4% | 5.7% | 5.0% | 2.9% | 5.2% | 5.6% | 5.2% | 4.4% |
| adm1 | 5.7% | 5.5% | 4.4% | 5.3% | 3.0% | 6.2% | 5.3% | 5.4% | 5.7% |
| adm2 | 4.8% | 4.7% | 3.9% | 4.1% | 2.3% | 5.1% | 6.2% | 5.0% | 0.0% |
| adm3 | 4.2% | 4.6% | 3.2% | 4.0% | 2.0% | 4.5% | 5.1% | 4.9% | 99.6% |
| **$p = 50$** |  |  |  |  |  |  |  |  |  |
| null | 3.1% | 3.6% | 3.7% | 4.0% | 2.0% | 3.4% | 3.5% | 3.8% | 4.4% |
| adm1 | 4.3% | 5.2% | 4.0% | 3.5% | 1.2% | 4.7% | 4.8% | 5.2% | 4.0% |
| adm2 | 4.1% | 4.4% | 4.8% | 4.7% | 2.4% | 4.0% | 5.3% | 4.9% | 0.0% |
| adm3 | 4.5% | 5.4% | 6.4% | 5.1% | 2.7% | 4.7% | 4.5% | 5.2% | 35.4% |

**Table 2.** Power estimates at a significance level of 5% for the FBAT, GTDT and RV-GDT statistics. We considered six scenarios, separately for *p* = 30 and *p* = 50 variants. All results based on 1,000 replicates. scenario 1: three causal variants, effect sizes 0.4| log_10_(*MAF*) |, same direction scenario 2: three causal variants, effect sizes 0.4| log_10_(*MAF*) |, different direction scenario 3: two causal variants, effect sizes 0.4| log_10_(*MAF*) |, same direction scenario 4: two causal variants, effect sizes 0.4| log_10_(*MAF*) |, different direction scenario 5: four very rare causal variants, effect size 1.0, same direction scenario 6: three causal variants, effect sizes 0.4| log_10_(*MAF*) |, same direction, in strong LD with other variants.

|  |  | FBAT |  |  |  |  | GTD |  |  | RV-GDT |
| --- | --- | --- | --- | --- | --- | --- | --- | --- | --- | --- |
|  |  | Burden | SKAT | MAX | HC | MAX-BF | GTD-AD | GTD-DOM | GTD-CH | RV-GDT |
| **$p = 30$** | 1 | 32.2% | 87.9% | 77.6% | 79.2% | 72.0% | 32.7% | 39.7% | 17.2% | 45.5% |
|  | 2 | 19.2% | 85.4% | 75.1% | 76.9% | 69.7% | 19.7% | 21.5% | 10.7% | 27.4% |
|  | 3 | 20.9% | 83.7% | 72.5% | 73.1% | 68.2% | 21.9% | 27.0% | 11.0% | 32.7% |
|  | 4 | 7.1% | 48.0% | 48.8% | 48.7% | 41.4% | 7.7% | 8.7% | 5.3% | 0.17% |
|  | 5 | 11.3% | 11.5% | 32.6% | 39.5% | 24.8% | 11.6% | 13.7% | 7.5% | 18.8% |
|  | 6 | 65.9% | 82.3% | 71.5% | 77.8% | 64.2% | 67.5% | 29.1% | 13.4% | 77.2% |
| **$p = 50$** | 1 | 50.2% | 90.8% | 77.8% | 80.9% | 67.9% | 51.3% | 31.8% | 17.9% | 64.1% |
|  | 2 | 24.0% | 88.3% | 73.5% | 75.3% | 64.9% | 25.2% | 12.1% | 9.9% | 35.6% |
|  | 3 | 32.9% | 83.4% | 68.6% | 70.0% | 61.3% | 33.4% | 18.9% | 11.7% | 47.6% |
|  | 4 | 8.2% | 50.6% | 43.3% | 45.8% | 34.3% | 8.7% | 4.9% | 6.4% | 0.7% |
|  | 5 | 8.3% | 11.0% | 26.8% | 31.4% | 17.9% | 8.5% | 9.0% | 7.6% | 13.5% |
|  | 6 | 72.6% | 84.9% | 69.3% | 78.5% | 58.0% | 73.7% | 21.9% | 13.5% | 83.3% |

In Table 1, we observe that all methods control the Type I error appropriately. The only exception is the RV-GDT in the scenario of population admixture with discordant parental phenotypes (adm2 and adm3, Table 1). This is expected, as the GDT/RV-GDT test compares the frequencies between affected and unaffected family members and cannot distinguish between association and admixture in the parents. We also note that the RV-GDT computes a one-sided p-value, which explains the deflation/inflation behavior, depending on the parental phenotypes. The power results in Table 2 demonstrate the advantages of non-burden tests in specific scenarios. The power results are also illustrated in Figures 1 and 2. The SKAT statistic shows the highest power in the first three scenarios and outperforms the other tests. However, the MAX and HC statistics also show substantial power. The results for scenario 4 are comparable between SKAT, MAX, and HC. In scenario 5, the HC statistics achieves the highest power, which is supported by our theoretical considerations as well (see Appendix A). In the last scenario 6, all tests achieve substantial power, as expected, due to the LD structure that pushes power. The most powerful tests here are SKAT and RV-GDT, but MAX and HC test statistics achieve similar results. If we have different effect directions (scenario 2 and 4), the burden test loses power compared to the consistent effect direction scenarios 1 and 3, which is expected. The FBAT Burden test and GDT-AD have almost no power in scenarios 4 and 5. It is important to note that the GTDT-AD and the FBAT burden test are essentially based on the same test statistic idea, the only difference lies in the fact that the GTDT assigns haplotypes (with possible error) and our approach uses the robust conditional genotype distribution computed by the FBAT haplotype algorithm.

**Figure 1.**
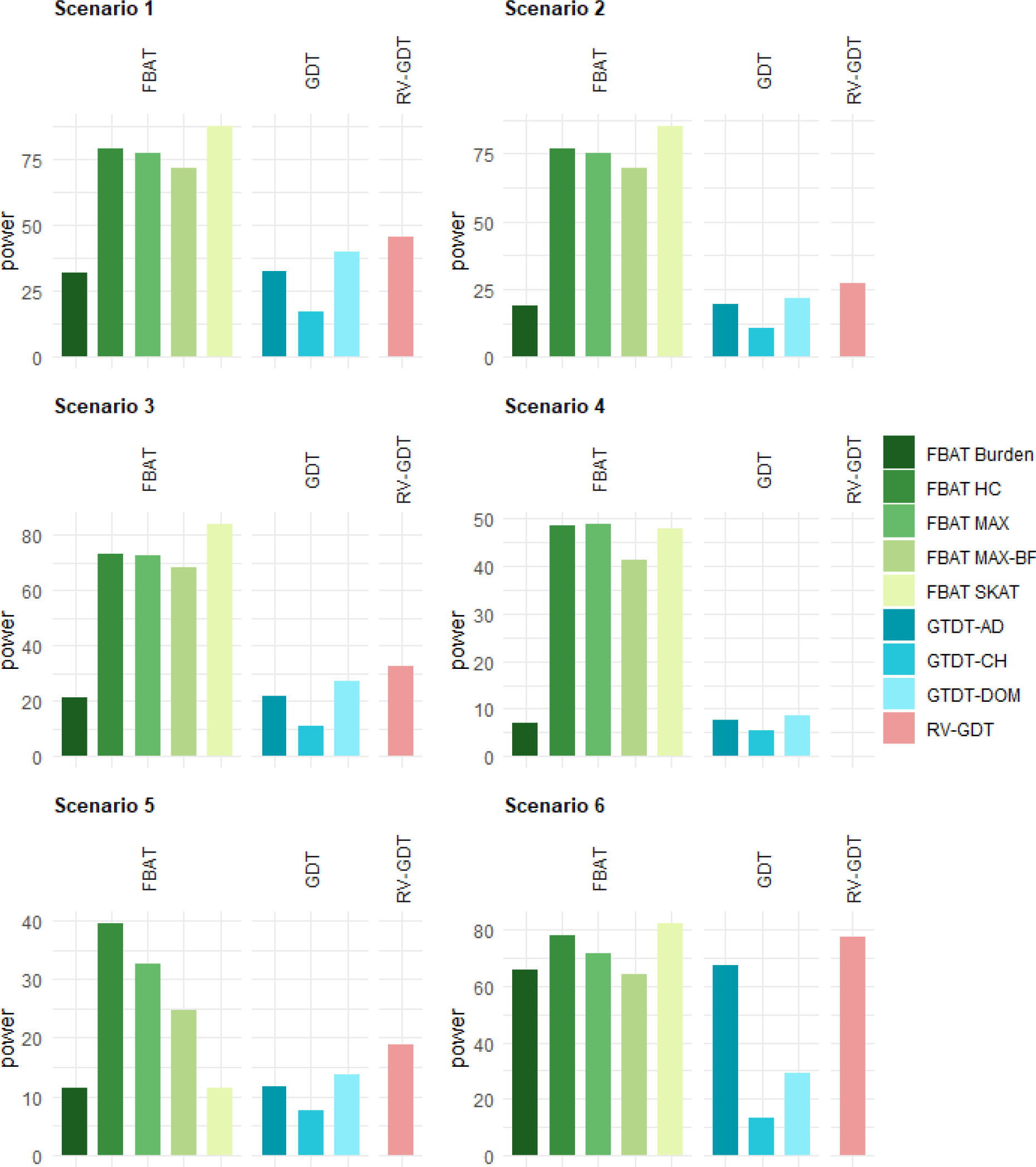
Power results in six different scenarios for genetic regions consisting of 30 variants at a significance level of *α* = 0.05. All results based on 1,000 replicates.

**Figure 2.**
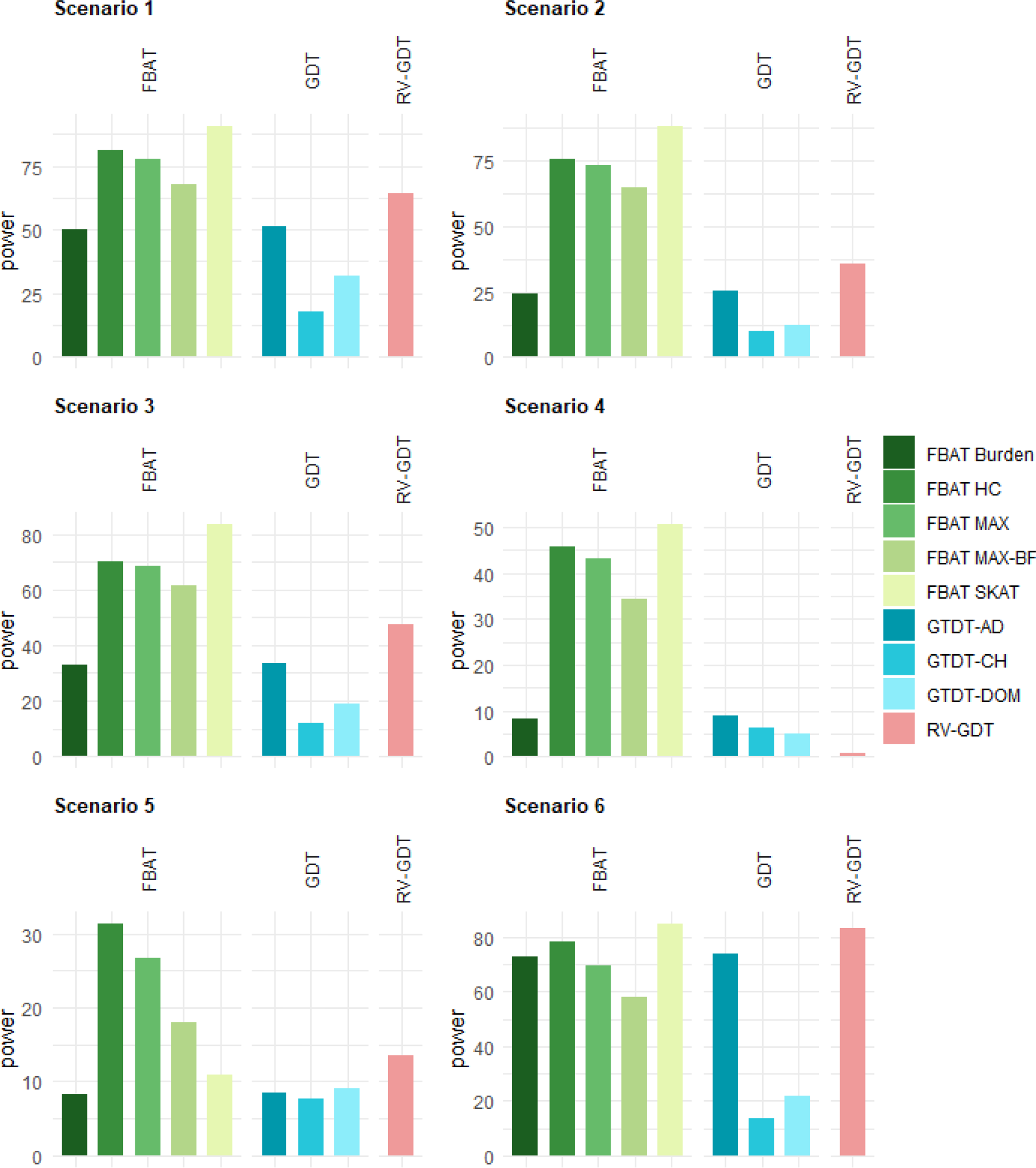
Power results in six different scenarios for genetic regions consisting of 50 variants at a significance level of *α* = 0.05. All results based on 1,000 replicates.

There is a substantial difference between the power of the MAX statistic and the MAX-BF statistic in all scenarios, because the Bonferroni correction is conservative, and the p-value of the MAX statistic is evaluated based on the joint conditional genotype distribution. We note that this difference could be even larger if the genome-wide significance levels for region- and single variant based testing are considered.

#### Dense genetic regions with phased data

For our second set of simulation studies, we utilized the 1006 EUR population haplotypes from the 1000 Genomes Project along 1,000 consecutive rare genetic variants with MAF below 3%. In this simulation study, we consider a large number of variants in combination with a sparse signal, which means a small subset of causal variants that are not in strong LD with any other variants. We simulated affected offspring trios as described in the first simulation study but also stored the phased haplotypes for all members of the trio. In the scenario where the haplotypes are observed, the conditional distribution identified by the FBAT haplotype algorithm equals the distribution where both parents transmit one of the observed haplotypes with equal probability of 0.5. We compared the performance of the FBAT, the GTDT, and the RV-TDT BRV ^27^ statistics to demonstrate the potential advantage of non-burden tests in the presence of sparse signals. Again, we also included the MAX-BF test, where we considered the Bonferroni corrected significance level based on *p* = 1,000 tests. As mentioned above, the knowledge about the phased haplotypes is the preferred setting for the GTDT. Also, the RV-TDT requires phased haplotypes ^27^. We considered a null hypothesis scenario and four different power scenarios (Table 3 and Figure 3). For the null hypothesis simulation and the first two power scenarios we simulated 1,000 trios; the last two power scenarios are based on 10,000 trios. The four power scenarios include causal variants that are in almost no LD with other variants, and the number of causal variants is small compared to the overall number of *p* = 1,000 variants. All results are based on 1,000 replicates, and the p-values for all test statistics were evaluated empirically based on the same 1,000 draws from the conditional haplotype distribution. The results for this simulation are also visualized in Figure 3.

**Table 3.** Type I error and power estimates at significance levels of 1% and 5%, all results based on 1,000 replicates. MAX-BF refers to the test that at least one single variant test statistic reached the Bonferroni-corrected significance level based on *p* =1,000 tests. scenario 1: two causal variants, MAF~0.2%, almost no LD with other variants, effect size 1.8 scenario 2: two causal variants, MAF~2%, almost no LD with other variants, effect size 0.7 scenario 3: 16 causal variants, MAF~0.1%, almost no LD with other variants, effect size 0.7 scenario 4: 16 causal variants, MAF~0.1%, almost no LD with other variants, effect size 0.7, effect direction alternates

|  |  | FBAT |  |  |  |  | GTDT |  |  | RV-TDT |
| --- | --- | --- | --- | --- | --- | --- | --- | --- | --- | --- |
|  |  | Burden | SKAT | MAX | HC | MAX-BF | GTDT-AD | GTDT-DOM | GTDT-CH | RV-TDT BRV |
| $\alpha = 0.05$ | null | 5.2% | 4.5% | 3.9% | 5.0% | 0.7% | 5.2% | 2.5% | 5.9% | 5.5% |
|  | 1 | 5.6% | 14.5% | 81.2% | 45.0% | 56.0% | 5.6% | 1.7% | 4.1% | 8.0% |
|  | 2 | 5.6% | 85.0% | 94.9% | 80.8% | 85.2% | 5.6% | 1.5% | 6% | 2.1% |
|  | 3 | 43.8% | 14.2% | 67.4% | 82.3% | 59.7% | 43.8% | 3.3% | 5.2% | 56.2% |
|  | 4 | 7.1% | 6.3% | 51.9% | 60.4% | 45.8% | 7.1% | 2.8% | 4.1% | 12.0% |
| $\alpha = 0.01$ | null | 1.3% | 1.5% | 0.7% | 1.2% | 0.0% | 1.3% | 0.4% | 1.1% | 1.3% |
|  | 1 | 1.3% | 3.4% | 61.2% | 39.9% | 33.0% | 1.3% | 0.2% | 0.8% | 2.3% |
|  | 2 | 0.9% | 54.6% | 87.6% | 77.7% | 71.0% | 0.9% | 0.2% | 1.3% | 0.5% |
|  | 3 | 23.0% | 2.7% | 39.2% | 48.3% | 32.0% | 23.0% | 0.5% | 1.3% | 32.2% |
|  | 4 | 1.8% | 0.9% | 27.3% | 31.9% | 20.8% | 1.8% | 0.5% | 0.8% | 3.1% |

**Figure 3.**
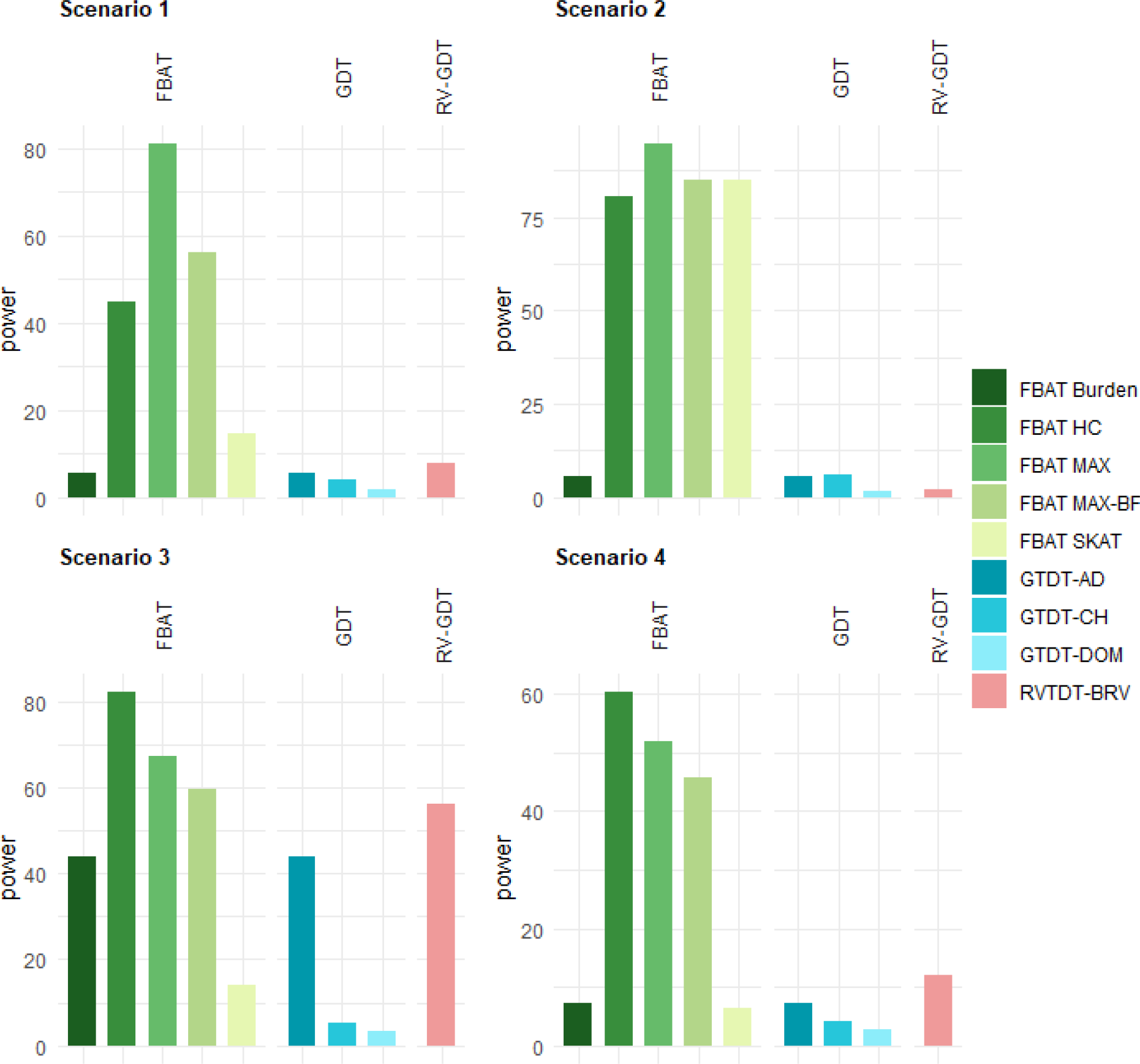
Power results for four different scenarios for genetic regions consisting of 1,000 variants at a significance level of *α* = 0.05. All results based on 1,000 replicates.

In Table 3, we observe that all test statistics control the Type I error appropriately. Since all p-values are evaluated empirically based on the conditional haplotype distribution by simulation, this is expected. In the first power scenario 1, the MAX test statistic achieves the highest power as we simulated a sparse and rare, but strong signal in the genetic region, consisting of two rare variants. The HC test statistic also achieves substantial power, whereas all other tests (except MAX-BF) are almost powerless in this scenario. In the second scenario, where the MAF of the two causal variants is much higher, the MAX test statistics still outperforms the other tests, but also the SKAT and the HC test statistics show good performances. In scenario 3, where many very rare causal variants have a relatively small effect size, the HC is the most powerful test. This is in line with the results in Mukherjee et al. ^20^ that describe a lower detection boundary in the mild sparse regime compared to the MAX test statistics. However, the RV-TDT BRV and MAX test statistic also achieve substantial power in this scenario. The power behavior differs more in the last scenario 4, where the effects are pointing in different directions. Here, the MAX and HC statistics have a significantly increased power compared to the other tests, while the HC test statistic is the most powerful one. We note that the FBAT burden and the GTDT-AD test are very close to the nominal level in scenarios 1, 2, and 4. Both tests are equivalent since the test statistics are the same.

Overall, again, the MAX test shows higher power than the MAX-BF tests, as described in the context of the first simulation study.

### Real data analysis

To demonstrate the applicability and the advantages of our proposed framework, we analyzed a whole-genome sequencing dataset consisting of 897 complete asthmatic trios from Costa Rica ^28^. After standard quality control, including Mendelian error rates, we excluded all variants with a MAF above 5%. The resulting 27,345,734 non-monomorphic variants were partitioned into approximately 547,000 consecutive windows of 50 rare variants. Other partitioning approaches could be considered here ^29,30^, but, as the focus of this data analysis was to demonstrate the feasibility of our approach, we did not explore different window-strategies here.

For each window, we performed the Burden, SKAT, MAX, and HC test, using affection status as the phenotype. We evaluated the p-values by simulation, where we used an adaptive heuristic that increases the number of simulations if the estimated p-value is close to the minimum possible value. The smallest possible p-value was *p* = 10^−9^, since the number of simulations was truncated at 10^9^. In Figure 4, we plotted the corresponding quantile-quantile-plot. The plot indicates that the test statistics control the Type I error rate but can also identify potential findings.

**Figure 4.**
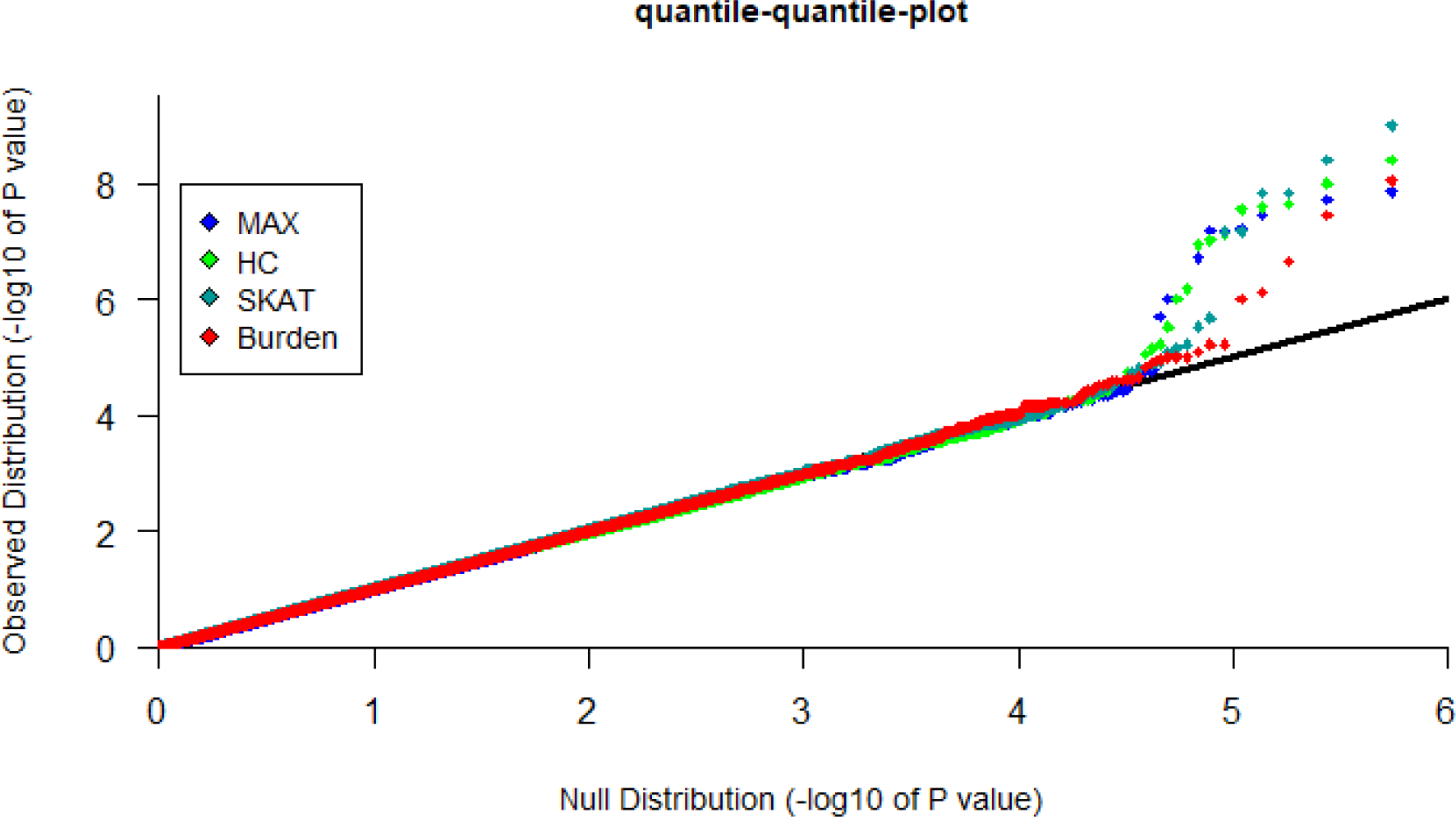
Real data analysis of 897 asthmatic offspring trios in a WGS study. Quantile-quantile plot for Burden, SKAT, MAX, and HC test statistics based on approximately 547,000 windows of 50 consecutive rare variants.

Based on the approximately 4 ∗ 547,000 tests and a False Discovery Rate at α = 0.05 ^31^, our approach identified three single significant regions on Chromosome 1, 12, and 21, as well as multiple consecutive significant regions on Chromosome 10. The significance of the three regions on Chromosomes 1, 12, and 21 was declared by the Burden test, whereas the single variant FBAT p-values within the regions were not in the range of genome-wide significance. The other regions on Chromosome 10 were identified by the MAX, HC, and SKAT tests. The lowest p-value of 10^−9^ was reached by the SKAT test. For all these regions, the Burden test did not reach the magnitude of genome-wide significance. This shows the benefits of combining different test statistics to identify distinct genetic signal structures.

## Discussion

In this manuscript, we propose a general framework for the region-based association analysis of sequencing datasets with family-based designs. The framework incorporates burden tests, SKAT, maximum and higher criticism approaches, and, given the flexibility of the framework, any future approach can straightforwardly be implemented. In contrast to previously published approaches, the joint genotype distribution along the loci is not obtained by empirical estimates, but via the haplotype algorithm for FBATs ^16,17^. This allows our proposed testing framework for FBATs to assess the significance of an arbitrary test statistic based on simulations or based on the exact distribution. This approach is enabled by the recent improvements in the FBAT haplotype algorithm ^16^, which reduces the computational burden of the original approach by several magnitudes. Our simulation results illustrate that the optimal test for the region-based analysis depends on the specific genetic architecture of the disease, and any WGS analysis relying on just one single test statistic may not detect all associations contained in the data. While dense signals with consistent effect directions can be captured by burden tests, different effect directions and less dense signals can be identified by SKAT approaches. If the signal becomes more separated and sparser, the MAX and HC approaches can be the most powerful tests.

The proposed implementation of the simulation-based p-values requires the user to pre-select the number of simulations that FBAT performs for each test. The computational burden can be decreased by adaptive strategies ^23^. The applications of the proposed analysis framework to simulated and real data illustrate that the theoretically expected advantages are also of practical relevance and that simulation-based p-values are not prohibitive in WGS settings. A subject of future research will be to integrate the existing FBAT approaches to multivariate phenotypes, longitudinal data, age at onset ^32,32–34^, gene-environmental interactions, and testing strategies into the proposed framework ^3,6,7^.

## Appendix A: Detection of sparse signals

We consider the scenario of an affected offspring trio. Both parental genotypes are observed along with the *p* variants in the analysis region. As noted in Chen et al. ^26^, if there is no variant where all three observed genotypes (mother, father, offspring) are heterozygous, the phase information can be recaptured from the observed unphased genotype data. However, as described in Hecker et al. ^14^, treating inferred haplotypes as observed haplotypes can lead to misspecification.

Nevertheless, more specifically, if there is no variant for which both parental genotypes are heterozygous, haplotypes can be phased, and the resulting conditional genotype distribution obtained by the FBAT haplotype algorithm equals the conditional distribution where we treat the haplotypes as observed. If we restrict the genetic data to rare variants, this is true for most nuclear families. In addition, with relatively high probability, at least 1 parent has only 1 minor allele in the genetic region.

Let us denote the phased parental mating type for such a trio by >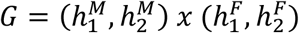 The possible offspring genotypes are denoted by >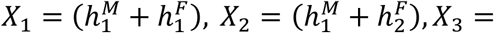 >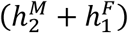 and >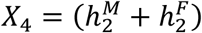. We assume the following, commonly used, disease model that describes the conditional offspring genotype distribution

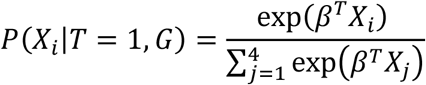

where the *p* dimensional vector *β* describes the genetic effects of the variants in the region.

If we denote the inherited offspring haplotypes by (*h*^*M*^, *h*^*F*^), this model factors into the product of the two likelihoods

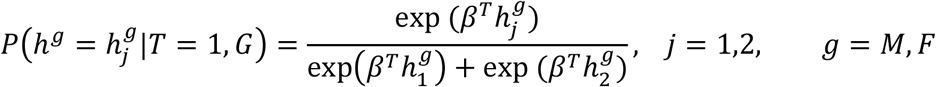

Since the haplotype data is sparse as described above, this setting matches the scenario of Weakly Correlated Designs that is described in the paper by Mukherjee et al. ^20^ about sparse binary regression (Definition 4.1). They showed that in the sparse regime, the higher criticism and the maximum statistic can identify sparse alternatives efficiently (see Theorem 7.4).

Although this motivates the application to affected offspring trios, the statistics can be applied in all FBAT scenarios. The Type I error is preserved because we utilize a simulation-based approach.

## Acknowledgments

This work was supported by Cure Alzheimer’s Fund; the National Human Genome Research Institute [R01HG008976]; and the National Heart, Lung, and Blood Institute [U01HL089856, U01HL089897, P01HL120839, P01HL132825].

## Web Resources

The FBAT software is available at https://sites.google.com/view/fbat-web-page. A new version that implements the described methodology is currently in preparation and will be available soon.

